# Vividness of mental imagery reflects a broad range of internally generated visual experiences

**DOI:** 10.1101/2025.09.30.679629

**Authors:** D. Samuel Schwarzkopf, Xinran A. Yu, Ecem Altan, Loren N. Bouyer, Blake W. Saurels, Elizabeth Pellicano, Derek H. Arnold

## Abstract

Research on mental visual imagery typically relies on vividness ratings. However, vividness is ill-defined as it lacks an objective reference. Here, we present survey results that suggest vividness is nevertheless a robust measure. It explains individual differences of a broad range of subjective experiences, from the detail of mental imagery, the propensity to report having other internally generated visual experiences, and the vividness of visual dreams. Critically, simple vividness ratings can replace the protracted questionnaires commonly used for this purpose and reduce methodological issues with these instruments. We further find that vividness is closely linked with the experience of “seeing” mental images or projecting them into the external world. People who report seeing mental images with their eyes shut are also more likely to experience externally projected imagery. Nevertheless, many people report having mental depictions but without seeing. Overall, our results indicate we should redefine visual aphantasia to distinguish individuals with faint or unseen visual images from those completely lacking a pictorial representation.

## Introduction

Mental imagery allows people to picture things inside their minds [1]. We might use imagery to recall events or to daydream. When reading a good novel, many people automatically picture the characters and scenes. Some people can close their eyes and picture themselves in a different place. On the flip side, mental imagery can intrude into our thoughts [2,3].

Many studies on mental visual imagery seem to have been designed in a way that assumes everyone who can visualise in fact *sees* their mental images in front of them [4–8]. In one study, participants were asked “to construct a mental image of [a] geometric shape” (such as a diamond) within a region on a test display marked by placeholders [7]. After this, a small dot would be presented on the test display, and participants were asked to decide if it had been presented inside or outside of their imagined shape. This implies that participants should be able to form a mental image that is coincident with a cued location on a physical test display. Such an experimental design does not seem to allow for the possibility that some people are able to visualise but unable to map their imagined experiences onto the physical world.

Another study asked participants to project an imagined grating onto a screen, to test how this would interfere with detecting an actual grating at that location [5]. The actual grating slowly faded in (see Figure 1A) “to mimic the gradual nature of mental image generation.” Not only does this imply that participants can see mental imagery as if it were in front of them in the external world, but also that it invariably emerges slowly and grows more intense over time. Similarly, in a study on how mental images interfere with a visual acuity task, participants were asked to match the apparent contrast of imagined black lines [8]. Rather than reporting they imagined black lines, all observers selected light grey lines, implying that they experienced their mental images as faintly blended into what their eyes see (Figure 1B).

**Figure 1.**
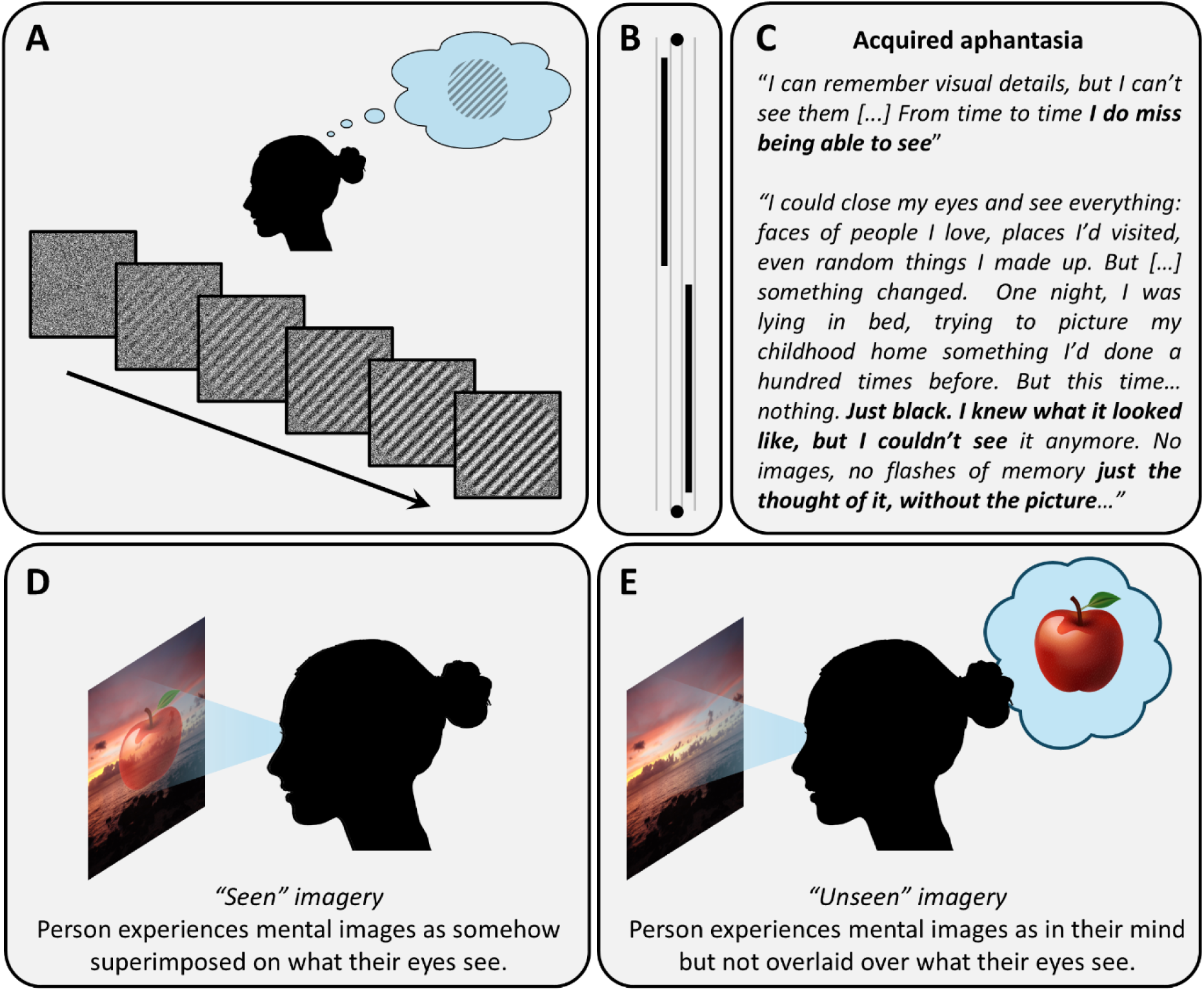
Do some people literally “see” mental imagery? The experimental design of some previous studies on imagery suggests they might. **A.** In one study [5], participants imagined an oriented grating while they viewed a white noise field. A real grating faded in on that stimulus in some trials “to mimic the gradual nature” of mental images. **B.** Another study [8] investigated how an imagined pattern of thin black vertical lines interfered with a Vernier acuity task (thick black line targets). For one condition, participants had to adjust the contrast of real lines to match the appearance of their imagined lines. All participants chose light grey lines (thin vertical lines in this image). **C.** Some cases of acquired aphantasia [9,10] describe the loss of mental imagery in terms of no longer seeing images, rather than losing the ability to have a picture in mind. **D.** Some people reportedly experience mental imagery as “seen,” superimposed on their visual input or even projected into the external world. A person with this experience will likely rate the vividness of the image by how opaque or saturated it appears before them. **E.** In contrast, a person who does not “see” imagery could still have a depiction of the object in mind – but judging the vividness is not straightforward. Unlike with seen imagery, there is no simple feature like opacity they can rate.

The experiences of individuals who have lost the ability to visualise (Figure 1C) are also illuminating in this context. One such individual with acquired aphantasia, MX, stated “I can remember visual details, but I can’t see them […] From time to time I do miss being able to see” [9]. Further, they reported that after some time their dreams “regained their visual qualities.” [9]. This implies that the visual quality of this person’s dreams was originally like his voluntary mental imagery. Another case of acquired aphantasia stated that when closing their eyes and attempting to “picture [their] childhood home” – something they used to do often – they could now only see “black” [10]. It follows from that description that before losing their mental imagery, they could indeed *see* something other than blackness when visualising their childhood home. Many of these descriptions are consistent with mental images being experienced as overlaid onto what is visible to our eyes, with imagined visual experiences being projected into our impressions of the external world (Figure 1D).

However, these described characteristics do not seem to match the subjective experiences of many people when they visualise. Rather, descriptions of “seeing” imagery can sound like hallucinations – but people typically feel that hallucinations and imagery (even if involuntary) are distinct. Evidently, given how many people have participated in such experiments without expressing bewilderment over the instructions, the experience of “seeing” imagery appears to be common. In a recent study asking people where they experienced their mental imagery, when people’s eyes were open nearly equal numbers (42 and 41%, respectively) reported that their imagery seemed to be located inside their head and in the external environment [11]. Moreover, 8% of people described their imagery as being located *somewhere else*. While these data can be interpreted in different ways, they highlight that the experience of mental imagery is more variable than many studies assume. For many people (including some of the authors), the experience of the mind’s eye may be more indirect, possibly even metaphorical, compared to the idea that imagery can intermix with our impressions of the external world (Figure 1E). We may consider our mental images to be purely inside our minds, somewhere off-screen [12], or at an undefinable location. It might be like having detailed *visual* knowledge without the sensation of *seeing*. When we close our eyes, we see black – and when we open our eyes, we do not experience any ethereal phantom images that can obscure our impressions of the external world. Nevertheless, we would consider our imagined experiences as pictorial. Could these differences be simply due to semantic disagreements in how different people describe the same subjective experience?

Imagery that is seen, especially when it is seen as if projected into the external world, could be a manifestation of hyperphantasia [13] or of eidetic visual memory [14]. One case study described an eidetic person who – when asked to recall a random dot pattern and “cast” it onto a display of a physical dot pattern – could experience the combined pattern as stereoscopic depth [15]. This person could also “hallucinate at will a beard on a beardless man, leaves on a barren tree, or a page of poetry in a known foreign language.” Importantly, they also reported that these “visions […] often obscure a real object.” This ability implies an intense and highly accurate spatial form of imagery that interferes with real visual inputs.

But seeing imagery need not dictate that mental imagery is necessarily strong or detailed. An image projected into the external world could appear transparent or schematic, and only barely obscure any real views. An image that is seen with closed eyes need not be photorealistic but could instead be experienced as a fuzzy shadow. Conversely, some people might consider their imagery to be intense and detailed, but still only see black when they shut their eyes. In short, seeing imagery and its vividness could be orthogonal dimensions.

Critically, consideration of these issues suggests that current measures of imagery, and definitions of its extremes (like aphantasia and hyperphantasia), could be confounded. The most popular traditional measures of mental imagery are the two versions of the Vividness of Mental Imagery Questionnaire (the VVIQ and VVIQ-2, respectively). These questionnaires invite people to attempt to visualise 16 (VVIQ) or 32 (VVIQ-2) different scenarios [16,17]. In the VVIQ-2, participants are also supposed to close their eyes while they generate their mental images, while in the VVIQ they are supposed to take the questionnaire once with open eyes and once with closed eyes. However, as far as we can tell, in recent decades many researchers have neglected these instructions. In each case participants must rate each scenario on how “vivid” their mental image was using a 5-point Likert scale. Failures to experience imagery (whatever that means) on a subset of scenarios would result in lower imagery “vividness” scores.

But vividness has no objective meaning. This is the classic Plato’s Cave conundrum: participants must use their own definition to decide if a mental image is vivid and they can only reliably reference these experiences against their own experiences – they do not have access to how imagery appears to other people. Consider visualising with closed eyes. If you see mental images in a similar way as you see visual dreams, you are likely to judge how intense these images appear. If this is how you have always experienced mental imagery, this should come naturally to you. By contrast, when you close your eyes and only see black – and you had this experience your entire life – you have no reason to assume that a vivid mental image should entail seeing intense pictures against your closed eyelids. You would be more likely to respond to the questionnaire in a way that rates the detail, complexity, or extent of the picture in your mind because this is all you know.

The same issue is even more profound for imagery experienced while the eyes are open. If you can indeed see your mental images before your eyes, rating vividness could be easy: you might base it on the clarity or opacity of the imagined image relative to what is behind it (Figure 1D). Vividness would simply be a measure of how strongly imagined images seem to obscure what your eyes see. But if you experience your mental images without literally seeing them, the situation is not as straightforward (Figure 1E). Such a person might even respond that they only “know” they are thinking of an object, because they cannot literally *see* it. That is the exact wording of the lowest point on popular imagery questionnaires [16,17]. Therefore, such a person may score low on vividness ratings due to confusion regarding what it means to *see* imagery – even though they might consider themselves as having mental imagery.

Researchers have suggested that mental visual imagery recruits the same neural substrates as actual visual perception [14,18–21]. Thus, imagery can sometimes interfere with perception [5,6,8,22,23]. Any such tendency could be greater for people who *see* imagery. Indeed, a recent study found some evidence to suggest that people who experience imagery as if it is in front of them, in the environment, experience greater imagery priming in a recognition task of physically seen words [22]. While this evidence was exploratory, it sounds a note of caution. By lumping together imagers irrespective of how they experience imagery, research findings based only on vividness ratings may be heterogeneous.

To understand imagery, and how it manifests in different people, we need greater understanding of how these factors – vividness, the sensation of seeing images, and the ability to project them externally – interact. Without a good understanding of these inter-relationships, we cannot be certain what most imagery studies have been measuring. Here, we report results of two online surveys, designed to capture how different people experience visual imagery. We incorporated self-reports on the vividness of people’s imagery, inspired by cartoon questionnaires we had encountered in internet discussions on this topic. Such cartoon instruments could provide a more sensitive way of measuring how intensely a person “sees” their visualisations, because they ask people to match their experience to an exemplar image (Figure 2). Moreover, we asked targeted questions to determine where people experienced their imagery, and tested scenarios that in most people seem to automatically evoke mental imagery (such as when people read evocative passages of fiction). We also included questions on other internally-generated visual experiences – like people’s dreams, daydreams, and hallucinations. This approach allowed us to examine how real life experiences of imagery (and conceptually similar phenomena) are related to traditional measures of imagery vividness [16,17]. In the first survey, we directly compared the answers to our novel questions to classic VVIQ-2 scores. We also included questions with free-text answers to test the contents of spontaneously generated descriptions of imagery. In the second survey, we adapted and replaced some of the questions. We also removed the VVIQ-2 and free-text answers to shorten the length of the survey, to encourage wider participation.

**Figure 2.**
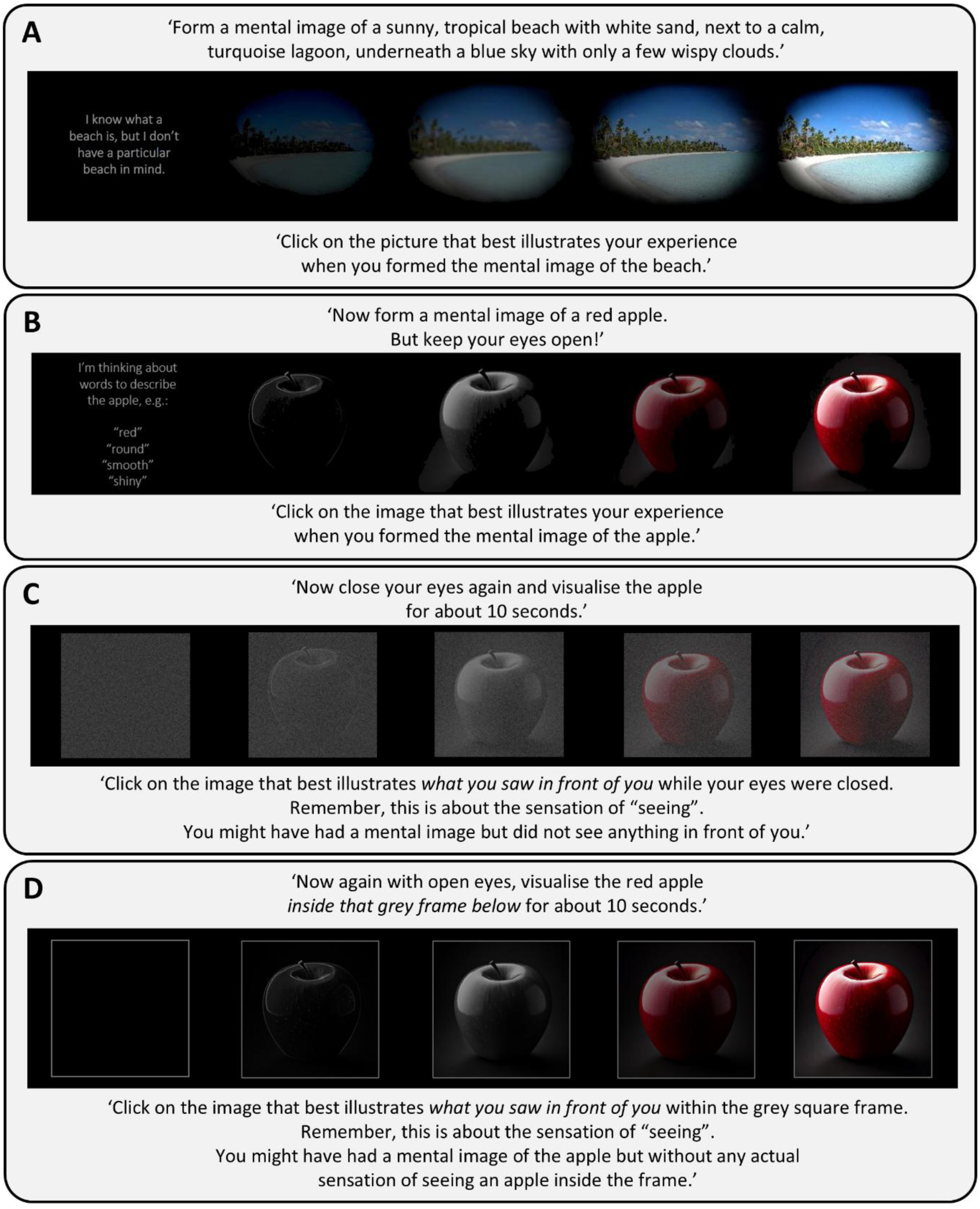
Questions probing mental imagery using a cartoon Likert-scale: the beach clarity (**A**), apple eyes-open (**B**), apple eyes-shut (**C**), and apple-in-frame (**D**) scenario. Both C and D explicitly instructed participants to judge the sensation of “seeing” mental images and clarified that they might have a mental image in mind without seeing anything.

## Materials and Methods

### Experiment 1

#### Participants

We recruited 100 participants through email and social media (Facebook, Twitter, Blue Sky) advertisements and via the Aphantasia Network (https://aphantasia.com). The survey was run online via the Testable platform (https://www.testable.org). We excluded 5 participants because they failed attention checks (see below), leaving 95 valid datasets. Participants’ mean age was 38.9 years (range: 16-80 years, two participants refused to answer), 39 identified as male, 52 as female, and 3 as non-binary (one participant refused to answer). All participants were shown information pages prior to the survey and were informed that by completing the survey they consented to their data being used for this research. All procedures were approved by the University of Auckland Human Participants Ethics Committee.

#### Questionnaires

The survey text was displayed in grey (RGB: 119, 119, 119) against a black background (RGB: 0, 0, 0). Most of the survey contained text questions, but there were several images. The full set of questions is available at https://osf.io/6r4hz. A demo version of the questionnaire is available at https://tstbl.co/458-340.

The survey started with questions about the propensity of various visual and/or mental experiences. These questions used a 5-point Likert scale (“Never”, “Rarely”, “Sometimes”, “Often”, and “Always”). We queried if they form mental images when reading fiction and how this interacts with movie/television depictions. We also surveyed them about visual experiences occurring without actual stimulation, such as dreams, near-sleep (hypnagogic or hypnopompic) visions, hallucinations after prolonged exposure to repetitive visual tasks (the “Tetris effect” [24]), and their experience of daydreams. Some questions led to forks in the questionnaire: e.g., if participants responded that they never dreamed, they would not be asked any further questions about dreaming and their responses to these questions were automatically coded as “Never.”

The next question asked which object was a darker green, a Christmas tree or a frozen pea, used previously as a paradigmatic example of a mental imagery task [25]. The following question asked participants *how* they answered this question, choosing between 1. Visualising the objects’ colours, 2. Simply knowing the answer, or 3. That they had guessed the answer. Another question instructed participants to think of a banana. They were then asked if the banana has black spots and given a range of response options, including Yes and No, two options of a mental image that does not allow them to answer this question, or that the question made no sense to them.

Participants also filled in the long-form VVIQ-2 [17] using the text from https://davidfmarks.net/vividness-of-visual-imagery-questionnaire-2. Each of the 32 questions were presented on a separate screen together with a slider that allowed for the five response options from the VVIQ-2. As per the original instructions, participants were asked to close their eyes each time they visualised a scenario.

The questionnaire continued with several questions probing how the participant experienced mental visual imagery. Specifically, we asked participants to form a mental image of a tropical beach, in whatever way they interpreted this instruction (beach clarity). We then presented them with five images of a tropical beach (One Foot Island, Aitutaki, Cook Islands) representing different levels of clarity, with the highest being a photorealistic depiction, and three versions with increasing blur, reduced saturation and brightness, with the lowest option consisting of a written description – implying that they had no pictorial sensation at all, but were merely aware of what features a beach has (Figure 2A). Another question (apple eyes-open) asked participants to imagine an apple and then presented them with four images of an apple at decreasing levels of clarity and saturation, plus a lowest option which only contained a written description of an apple (Figure 2B). This question explicitly instructed participants to keep their eyes open. The scaling of the visual features of the apple was not identical to what we had used for the beach scene. In both cases, the scaling of blur, saturation, and brightness was not linear but rather aimed to include levels of detail and “vividness” ranging from no visual imagery to highly photorealistic imagery.

The apple eyes-open question was followed by a question in which we showed four cartoons (in randomised locations) and asked which one of these best represented how they experienced their mental image of the apple (Figure 3A): an apple in front of the head (“Projector”), an apple inside the head (“Insider”), an apple in a thought bubble (“Off-screener”), and a thought bubble containing only adjectives describing the apple (“Verbaliser”).

**Figure 3.**
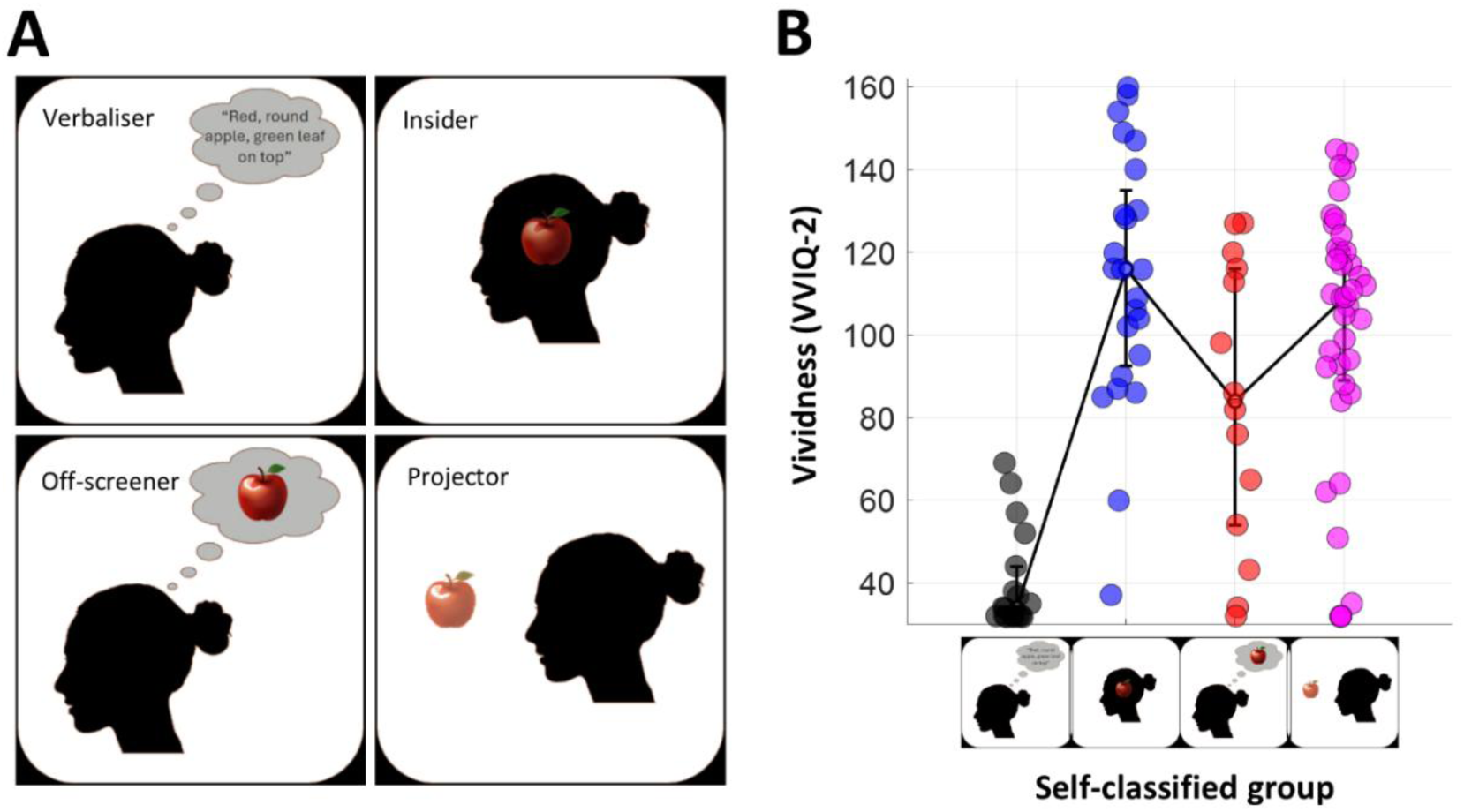
Results of Experiment 1. **A.** Cartoon depictions used for self-classification into the four groups. **B.** Vividness (VVIQ-2) scores for each group. Each dot is one participant. Some Gaussian jitter was added to dot locations for visualisation. The solid black lines connect the medians (error bars denote the interquartile range). Black: Verbalisers. Blue: Insiders. Red: Off-screeners. Purple: Projectors.

Next, to probe if participants literally see their imagery, we asked them to visualise an apple while their eyes were closed (apple eyes-shut). We then presented them with five cartoons depicting how the apple had appeared against the darkness (eigengrau) of their shut eyelids (Figure 2C). These ranged from a speckled dark noise field that aimed to evoke the impression of “eigengrau” or “eigenlicht” [26] to a photorealistic apple superimposed on that noise field. We included the eigengrau field to avoid participants mistaking the lowest option for the complete absence of mental imagery. It was instead intended to imply that this was about literally “seeing” an image against closed eyelids.

We then repeated this exercise with open eyes while presenting a grey square outline on the screen and instructed participants to visualise the apple inside the frame (apple-in-frame). We presented them with cartoons ranging from an empty frame to a frame containing a photorealistic apple (Figure 2D). Importantly, for both the apple eyes-shut and apple-in-frame scenarios we instructed participants to answer based on the sensation of “seeing” the apple, not merely if they had a mental image in mind. Cartoon questions were coded on a Likert scale of 1-5.

We also posed two questions allowing free-form answers. These questions asked participants to “think of a triangle” and to “think of a pink elephant,” respectively. Then a text entry box appeared, and participants were asked to describe the object they had thought of. Two questions in the survey were attention checks. These instructed participants to choose a particular option. We excluded five participants who gave an incorrect answer to one or more of these questions. Finally, participants had the option to report their age and gender but were informed that this information was non-mandatory.

#### Statistical analyses

Using the self-classification question, we divided participants into four groups based on how they experienced their mental image in the first apple question, that is into Projectors, Off-screeners, Insiders, or Verbalisers. We compared any questionnaire scores with Likert scales (including the summed VVIQ-2) between these groups using the Kruskal-Wallis test and Mann-Whitney U-test for pairwise comparisons. Despite the number of comparisons, we did not correct these for multiple comparisons, as this was an exploratory set of analyses aimed at revealing potentially subtle differences between self-classified groups.

We further compared all questionnaire items with numeric 5-point Likert scores. This included all the questions on mental experiences described above (e.g., propensity for daydreams, or of experiencing the Tetris effect), and the cartoon scale questions (beach clarity, apple eyes-open, apple eyes-shut, apple-in-frame). We also compared these variables to VVIQ-2 scores (which are themselves the sum of 32 5-point Likert items). To this end, we calculated a correlation matrix between all pairwise comparisons between all these variables, using the non-parametric Spearman’s ρ correlation (Supplementary Figure S1A). As this entailed 253 comparisons, we Bonferroni corrected p-values for these tests.

#### Analysis of free-text data

Three authors (DSS, AXY, EA) rated the free-text answers to the triangle and pink elephant questions for each participant on a 5-point scale, in terms of their detail and richness – with 1 being no imagery and 5 being the highest, most evocative descriptions. All three raters self-reported as having mental imagery, albeit of different manifestations (2 Offscreeners, 1 Projector). Essentially, these authors rated how clear and detailed a mental image was evoked by the participants’ descriptions. Ratings showed close agreement for both the pink elephant (Cronbach’s α=0.89) and the triangle questions (α=0.94).

### Experiment 2

#### Participants

We recruited 253 participants through email and social media (Facebook, Twitter, Blue Sky) advertisements and on Testable Minds. The survey was run online via the Testable platform (https://www.testable.org). We excluded 23 participants because they failed an attention check item (see below), leaving 230 valid datasets. Participants’ mean age was 31.1 years (range: 18-88 years, 4 participants refused to answer), 115 identified as male, 108 as female, and 4 as non-binary (3 participants refused to answer). All participants were shown information pages prior to the survey and were informed that by completing the survey they consented to their data being used for this research. All procedures were approved by the University of Auckland Human Participants Ethics Committee.

#### Questionnaire

Most methods were the same and several questions were identical to Experiment 1. The full set of questions is available at https://osf.io/8k2f5. A demo version of the questionnaire is available at https://tstbl.co/407-017.

We removed the VVIQ-2 questionnaire; and instead used the beach clarity question as a proxy measure of imagery vividness (Figure 5A). We also removed questions that did not probe visual experiences. Specifically, we only asked if participants have visual dreams or visual daydreams. We added a clarification to the question about whether participants imagined fiction –that this was about their mind’s eye (however they interpreted that), and that this should not necessarily imply that they “literally” see what they have imagined. Conversely, we asked explicit questions about how often they literally saw their daydreams, or experienced mental images as floating before them when they visualised objects or scenes. Instead of asking about the vividness of visual dreams, we only asked about the frequency of visual dreams.

We removed questions with free-text answers as well as the banana question. Instead, one question instructed participants to think of a triangle and then asked them if they knew what colour the triangle was. They could choose between three options: 1. Yes, 2. No, or 3. This question makes no sense. Another question asked them what kind of thinker they are, making them choose between 1. Being a visual thinker, 2. Hearing their inner voice but no visual thinking, 3. Thinking in words but without an inner voice, or 4. None of the above.

Finally, to probe how participants interpreted the phenomenon of aphantasia, we asked them what happens when an aphantasic person thinks of an object. The options were 1. “They only know what the object looks like, but they don’t literally see it”, 2. “They have no picture in their mind at all, but instead think in words, facts, or concepts”, or 3. “I don’t know - I’ve never heard of aphantasia.”

We also included one question as an attention check. This presented participants with pictures of apples from one of the imagery questions, but in reverse order. We instructed people to choose the second picture from the left. We excluded all participants who failed to respond correctly to this question.

Finally, we had initially included two questions to further probe the propensity of people to experience internal imagery. However, after starting the survey we realised that the phrasing of these questions contained a double negative, rendering responses ambiguous. We therefore removed these questions and data already collected from these questions were not analysed.

## Results

### Experiment 1

Based on their self-classification (Figure 3A), 41.1% participants in Experiment 1 reported the cartoon of an apple in front of their head best depicted their mental imagery experience (“Projectors”), 25.3% chose the option with the apple inside the head (“Insiders”), and 14.7% chose the option with the apple in a thought bubble (“Off-screeners”). Henceforth, we will refer to these three groups collectively as imagery groups. In contrast, those who chose the option that described the apple with a list of adjectives (“Verbalisers”) comprised 18.9% of our sample. Note that these proportions should not be taken as indicative of distributions across the general population, as participants for this study were recruited via advertising on websites that targeted people who self-report as having aphantasia.

Vividness ratings (VVIQ-2 scores) from our sample covered the entire possible range, with 12 participants scoring the minimum of 32 and one scoring the maximum of 160. There are no established criteria for what constitutes high or low vividness scores, and operational definitions of this are often arbitrary. Some studies define any participant as aphantasic when their equivalent VVIQ-2 scores are as high as 64 [1,27–30]. We do not aim to draw conclusions on imagery extremes solely based on vividness scores. However, for reference to other studies – 21% of our sample had vividness scores below 40 and 3% scored above 152. These thresholds correspond to participants who respond almost exclusively with the lowest and highest options in the VVIQ-2.

Importantly, the four groups based on self-classification (Figure 3B) differed significantly in their VVIQ-2 scores (Kruskal-Wallis test, χ^2^=36.9, p<10^−7^, uncorrected). Verbalisers had less vivid imagery than the other groups (Mann-Whitney U-test, all p<0.001). Off-screeners scored lower than Insiders (p=0.0142) but not Projectors (p=0.1178), and Insiders and Projectors were not significantly different (p=0.2234). Notably, the distributions of scores for the three imagery groups were long-tailed: each of these groups contained individuals with low VVIQ-2 scores, some below 40.

#### Correlations with vividness

Vividness scores correlated with many numerical measures from questionnaires (Supplementary Figure S1A; also see Supplementary Information for further discussion). Unsurprisingly, vividness ratings correlated strongly with other general vividness questions, especially with choosing how clear the imagined beach scene was (Figures 4A). Given this strong association, it is unsurprising that the four groups from self-classification show the same pattern as they did for VVIQ-2 scores (Supplementary Figure S2).

**Figure 4.**
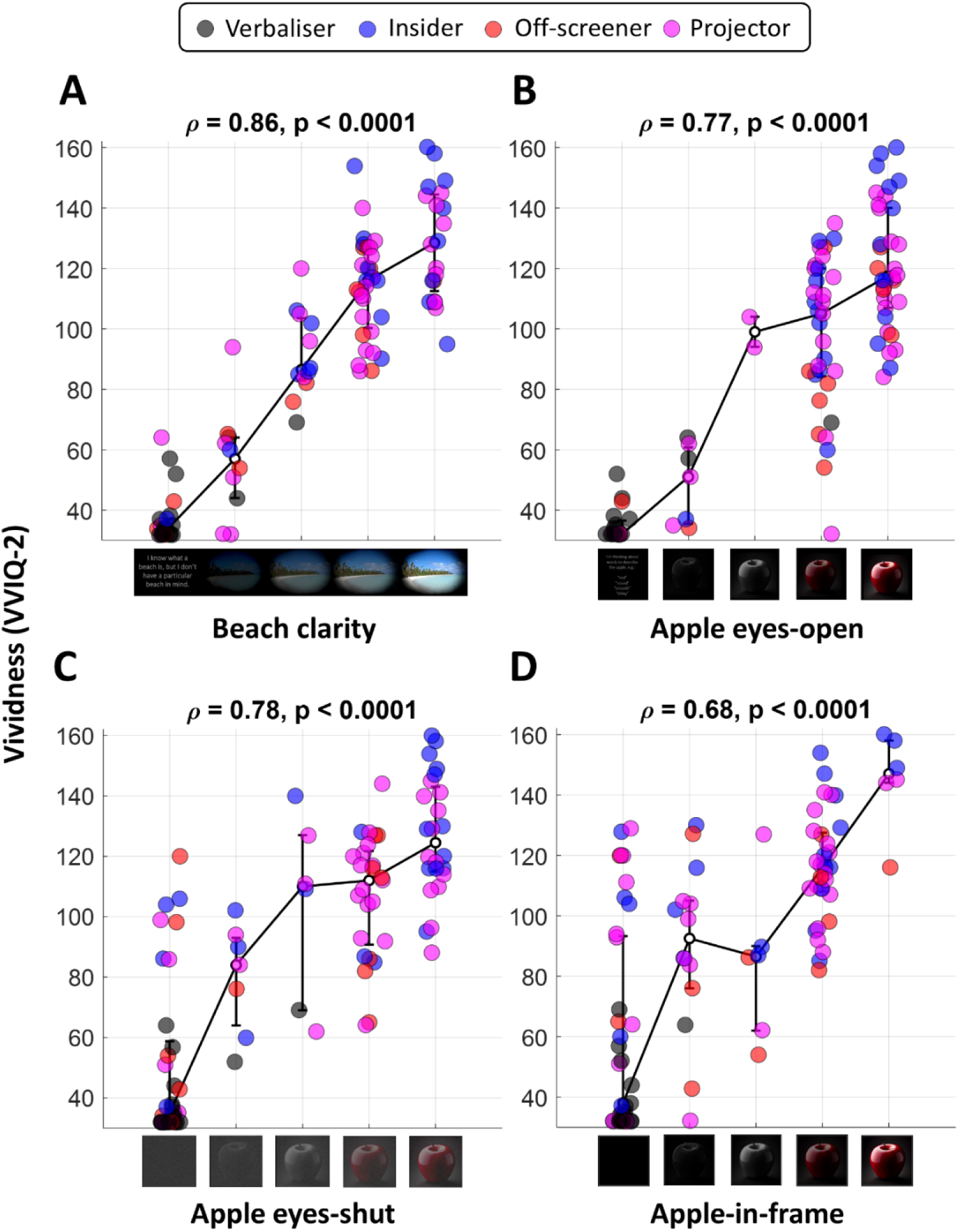
Vividness (VVIQ-2) scores plotted against the Likert score on the beach clarity (**A**), apple eyes-open (**B**), apple eyes-shut (**C**), apple-in-frame (**D**) scenarios. Each dot is one participant. Some Gaussian jitter was added to dot locations for visualisation. The solid black lines connect the medians (error bars denote the interquartile range). Black: Verbalisers. Blue: Insiders. Red: Off-screeners. Purple: Projectors. Statistics at the top of each panel show the Spearman correlation (Bonferroni corrected).

Vividness also correlated strongly with how an apple appeared in the mind’s eye (Figures 4B). It further associated with whether participants typically imagine scenes and characters when reading fiction (Spearman’s ρ=0.74, p<0.0001, Bonferroni corrected) and with how often people reported experiencing their daydreams visually (ρ=0.72, p<0.0001). It further correlated with how often people reported that movie or television depictions clash with their visualisations of the same characters, scenes, or events when reading (ρ=0.54, p<0.0001).

Vividness also correlated with other internally-generated experiences, such as the intensity/opacity of visual dreams (ρ=0.52, p=0<0.0001), the frequency of near-sleep (i.e., hypnagogic/hypnopompic) visions (ρ=0.53, p<0.0001), or the “Tetris effect” [24], and hallucinations reported after performing repetitive tasks (ρ=0.59, p<0.0001).

Critically, vividness also correlated with seeing the mental image of an apple against the black of closed eyelids (Figure 4C) or as projected within a frame on the screen with open eyes (Figure 4D). Scores for these two questions also strongly correlated with each other (ρ=0.69, p<0.0001). However, in both scenarios many participants with generally vivid imagery indicated that they did not see images at all: 15.2% in the apple eyes-shut and 21.6% in the apple-in-frame scenario scored in the top half of the vividness spectrum (VVIQ-2>96). Notably, such individuals did not consistently identify as Insiders or Off-screeners but also as Projectors (see upper-left corners of Figures 4C,D). Vividness scores also correlated with a tendency to experience daydreams as projected into the environment (ρ=0.61, p<0.0001), though again many individuals (25.5%) in the top half of the vividness spectrum responded that they never see daydreams projected externally. These correlations were not obviously linked to the self-classification of different people into different groups: individuals in the three imagery groups responded across the scale – only Verbalisers overwhelmingly chose the lowest options (no image/never).

Verbalisers were not the sole drivers of this association. When we repeated these correlational analyses only on imagery groups (i.e., without Verbalisers), we nevertheless observed strong and significant correlations between VVIQ-2 scores and these questions (Supplementary Figure S3). Especially the apple-in-frame question (Supplementary Figure S3D) showed that many participants who reported not seeing anything in the frame had average or even relatively high VVIQ-2 scores (median: 79, range: 32-129). This stands in stark contrast to the median VVIQ-2 of 38 when including Verbalisers.

#### Seeing imagery

Next, we analysed the propensity of people to report seeing mental images before their eyes. In both relevant apple scenarios (Figure 2C,D), scores of the four groups of people differed significantly from each other (both p<0.0001). Unsurprisingly, this was partly driven by Verbalisers, whose scores were lower than those of the three imagery groups (all p<0.003). In fact, Verbalisers overwhelmingly chose the lowest option, indicating that they did not see any image. However, at least for the eyes-shut scenario, Off-screeners also scored lower than Insiders (p=0.0165) and Projectors (p=0.023), while the latter two groups were not significantly different (p=0.5056). For the apple-in-frame scenario, the three imagery groups did not differ (all p>0.3461).

The four groups differed also in their tendency to experience daydreams projected into their visual field (χ^2^=28.4, p<10^−5^). Again, Verbalisers scored significantly lower on this item than the other groups (all p<0.0075), because all individuals in this group reported never seeing their daydreams projected (this includes individuals who responded they never daydreamed at all). Off-screeners also scored significantly lower on this than Insiders (p=0.0383) or Projectors (p=0.0021). The latter two groups were not significantly different (p=0.3255).

#### Colour comparison

A few questions did not use the 5-point Likert scale. They probed how participants typically use imagery. In one question, we asked participants to respond which object was a darker green, a Christmas tree or a frozen pea [25]. Participants overwhelmingly responded that the Christmas tree was darker; only two individuals chose the frozen pea. Curiously, both identified as Projectors, with VVIQ-2 scores of 32 and 109, respectively. Asked *how* they judged the object’s colour, 70.5% of participants thought they had visualised the objects and compared their colours. By contrast, 25.3% thought they simply “knew” the answer, and 4.2% reported having guessed. Vividness scores were significantly different between these three cohorts (χ^2^=23.4, p<10^−5^). Individuals who reported visualising had significantly higher VVIQ-2 scores (median: 110) than those who knew (median: 36, p<10^−7^) or guessed (median: 32, p=0.0012).

The four self-classified groups also differed on this question (χ^2^=21.8, p<0.0001). Verbalisers tended to respond that they knew the answer compared to Insiders and Projectors (both p<0.0001). Verbalisers and Off-screeners were not significantly different (p=0.0638). Focusing only on Verbalisers, 61.1% reported knowing the answer and only 11.1% reported that they had guessed. Interestingly, this leaves 27.8% who said they visualised the objects. The other groups overwhelmingly reported having visualised the objects, especially Insiders (87.5%) and Projectors (82.1%), but also most Off-screeners (64.3%). No Insiders and only a handful of the others reported guessing (Off-screeners: 7.1%, Projectors: 2.6%).

#### Contents of evoked imagery

Three questions tested whether thinking of an object automatically evokes mental visual imagery. First, we asked participants to think of a banana and then to choose one of five options that best described their experience. Two options were that the banana was either all yellow or that it had black spots; 23.2% and 45.3%, respectively, chose these options. Individuals choosing either answer had similar median VVIQ-2 scores (all-yellow: 114.5, black-spots: 112). Of the remaining participants, 7.4% responded that the image of the banana was too unclear to tell if it had black spots, and 7.4% responded that they could only tell the banana’s shape. Finally, 16.9% responded that the question made no sense to them.

Given both the all-yellow and black-spots answers were chosen by individuals with similar vividness scores, we pooled those two responses and recoded them as the maximum on a 4-point scale. The four self-classified groups differed significantly on this score (χ^2^=38.9, p<10^−7^). The three imagery groups did not differ significantly from one another (all p>0.1045), as most chose options with colourful imagery (Insiders: 91.7%, Off-screeners: 71.4%, Projectors: 82.1%). Some off-screeners (21.4%) responded that the question made no sense to them, while 7.1% found their image too vague, and nobody reported only knowing the banana’s shape. Only a few Insiders (4.2%) reported only knowing the shape, none reported having a vague image, and for 4.2% the question made no sense. Finally, 5.1% of Projectors reported having a vague image, 2.6% could only tell the banana’s shape, and 10.3% thought the question made no sense.

These results stand in stark contrast to Verbalisers, who were more likely to endorse weak imagery options than the other groups (all p<0.0028). Indeed, only one Verbaliser (5.6%) responded that the banana was all yellow. A large proportion of this group (44.4%) thought that the question made no sense, 27.8% reported only being able to tell the banana’s shape, and 22.2% thought that the banana was too vague to answer the question.

Two questions allowed free-text answers: participants were asked to think of a pink elephant and a triangle, respectively, and to describe their imagined experiences in words. Vividness scores predicted our ratings of the detail-richness of free-text answers. The answers by participants with higher VVIQ-2 scores tended to contain richer, more visually evocative detail. This was especially the case for the pink elephant (ρ=0.67, p<0.0001), while the association was less clear for the triangle (ρ=0.46, p=0.0005). A qualitative evaluation of these free-text answers also supports this (Supplementary Information).

#### Internal consistency of vividness scores

As reported, vividness, quantified by VVIQ-2 scores, correlated strongly (ρ=0.86) with our 5-point rating scale for the beach clarity scenario. The subdivision into the four groups from self-classification also aligned closely with VVIQ-2 (Figure 3B) and beach clarity scores (Supplementary Figure S2). Indeed, the correlations for either score with all the other questionnaire items were very similar (compare first two columns of the correlation matrix in Supplementary Figure S1A). This suggests that the simple beach clarity scale conveyed largely the same information as the protracted vividness questionnaire.

The VVIQ-2 contains 32 mental imagery instructions, grouped into eight scenarios. Participants must respond with the same 5-point rating scale to each item. For participants who do not vary much in their responses, this task can quickly become tedious. It could also entrain responses biases, especially with people at the extremes of the vividness scale. Self-reported aphantasics and Verbalisers may just repeatedly choose the lowest point without carefully attending to the specifics of item instructions.

To test this, we calculated the correlation matrix between all 32 items in the VVIQ-2, plus our two general vividness measures, the beach clarity and apple eyes-open scenarios. With 34 variables, this results in 561 unique pairwise correlations. Unsurprisingly, all comparisons were strongly correlated (mean ρ=0.78) with 95% of these correlations ranging between 0.68 and 0.87 (Figure 5A). Thus, both our general imagery scenarios correlated better with overall VVIQ-2 scores than most items in the VVIQ-2 correlated with one another. This suggests that the VVIQ-2 is at least somewhat redundant.

**Figure 5.**
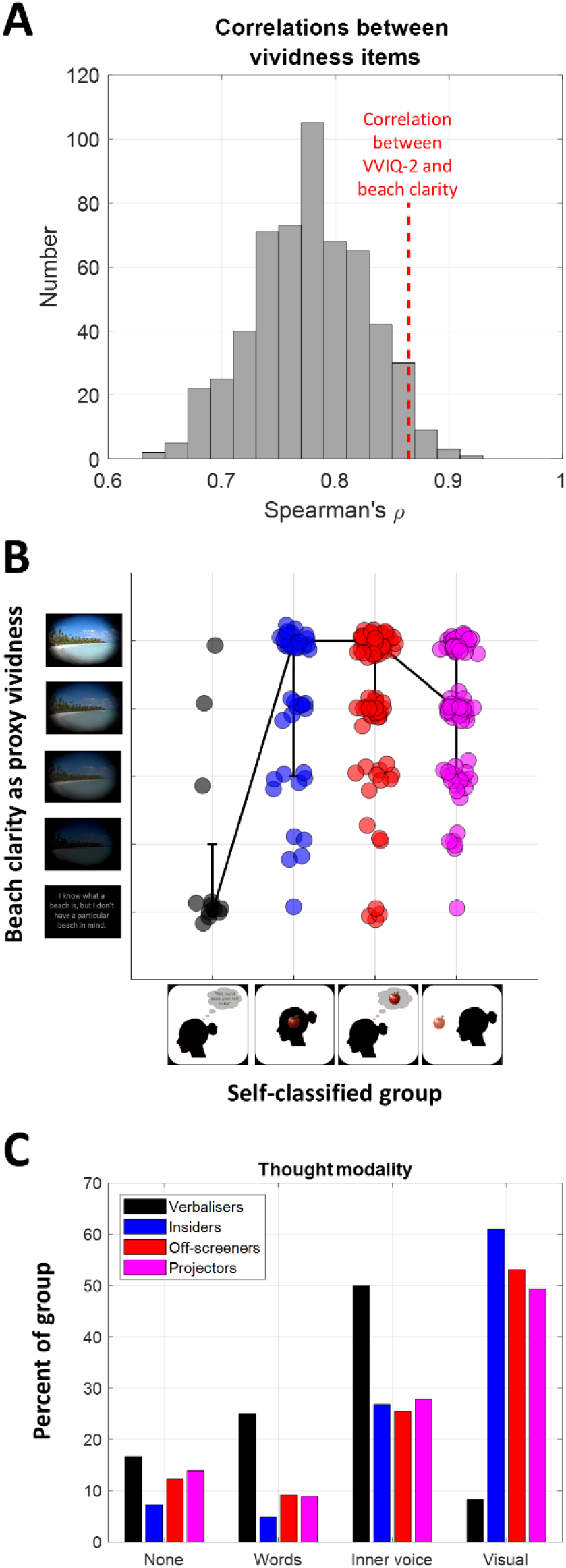
Beach clarity score as proxy vividness in Experiment 2. **A.** Histogram of all pairwise correlations (Spearman’s ρ) between individual vividness items in Experiment 1, including all 32 scenarios in the VVIQ-2, and our beach clarity and apple eyes-open scenarios. The red dashed line indicates the correlation between overall VVIQ-2 scores and the beach clarity scenario. **B.** Proxy Vividness (beach clarity) scores for each group from self-classification. Each dot is one participant. Some Gaussian jitter was added to dot locations for visualisation. The solid black lines connect the medians (error bars denote the interquartile range). **C.** Frequency histogram of members in each group who endorsed a given answer on the thought modality question. Black: Verbalisers. Blue: Insiders. Red: Off-screeners. Purple: Projectors.

### Experiment 2

We ran a second survey to probe some of these relationships in more detail, and to test the replicability of these findings. Due to the redundancy of the VVIQ-2 items, we decided to omit this questionnaire and instead we used the beach clarity question as a proxy measure of vividness. This shortened the length of the survey and allowed us to recruit a larger sample. Moreover, it should also reduce problems with entraining response biases in individuals who respond the same to each VVIQ-2 item.

Moreover, Experiment 1 had recruited from online communities interested in imagery extremes. Our data suggests that this caused a sampling bias by recruiting an unrepresentatively large number of potentially aphantasic individuals. In Experiment 2, we therefore decided to recruit more broadly.

#### Self-classification into groups

Based on their self-classification, 34.3% of participants were Projectors, 17.8% were Insiders, and 42.6% were Off-screeners (Figure 5B). Only a small percentage (5.2%) identified as Verbalisers. Conceptually replicating Experiment 1, the four groups differed in proxy vividness (χ^2^=32.3, p<10^−6^). This was driven solely by Verbalisers whose scores were lower than the imagery groups (all p<0.0001), although three Verbalisers responded with comparably high ratings (score ≥3). Off-screeners scored vividness *higher* than Projectors (p=0.001). The other groups were not significantly different (both p>0.2037). Notably, as in Experiment 1, all three imagery groups contained individuals who rated their vividness low.

#### Correlations with vividness

Proxy vividness scores correlated strongly with the intensity/clarity of the imagined apple (Figures 6, see also Supplementary Figure S1B for full results). However, similar to Experiment 1, more than half of participants (apple eyes-shut: 51.4%, apple-in-frame: 52.8%) who scored high on proxy vividness (beach clarity rated 4 or 5) scored low on the strength of the apple image seen with eyes shut (Figure 6B) or as projected inside the frame on the screen with open eyes (Figure 6C).

**Figure 6.**
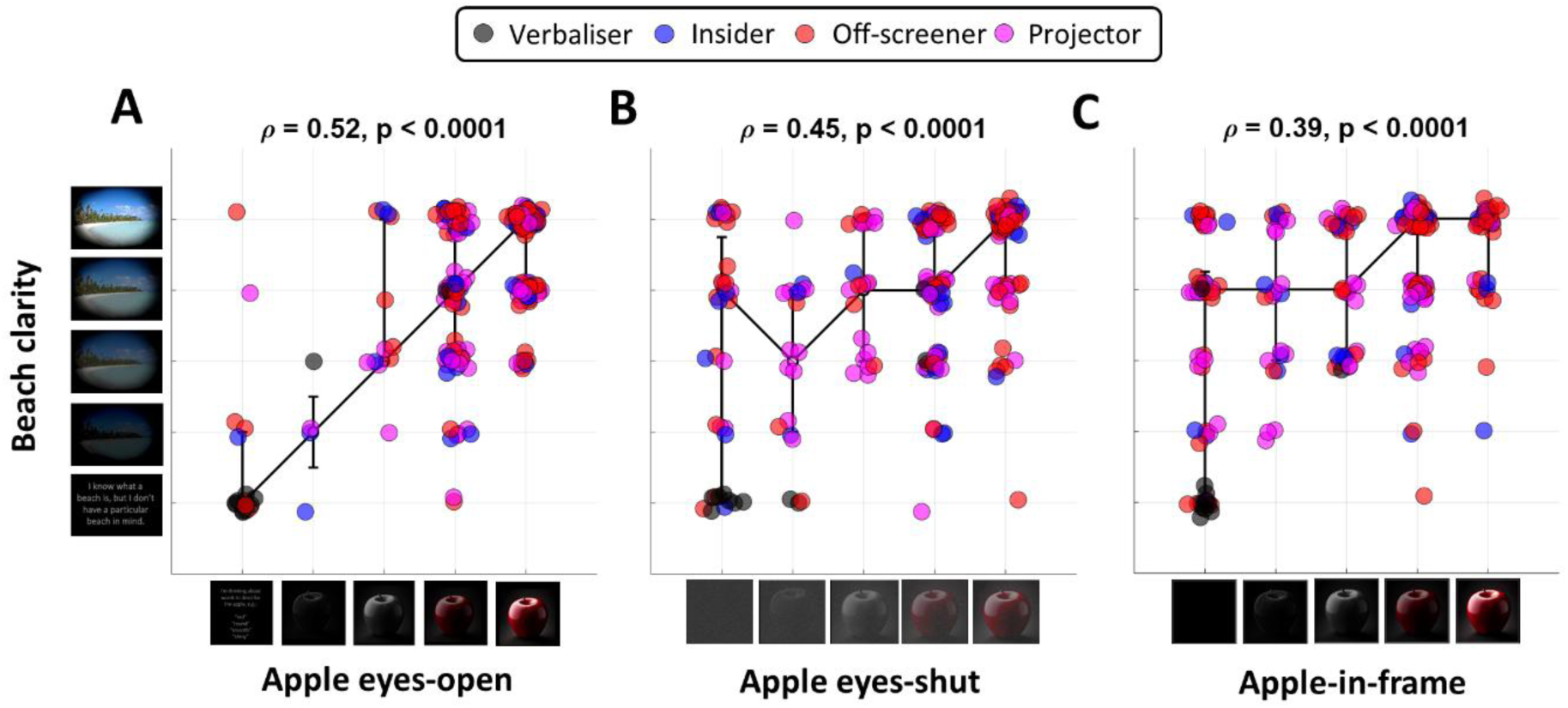
Proxy vividness (beach clarity) scores plotted against the Likert score on the apple eyes-open (**A**), apple eyes-shut (**B**), apple-in-frame (**C**) scenarios. Each dot is one participant. Some Gaussian jitter was added to dot locations for visualisation. The solid black lines connect the medians (error bars denote the interquartile range). Black: Verbalisers. Blue: Insiders. Red: Off-screeners. Purple: Projectors. Statistics at the top of each panel show the Spearman correlation (Bonferroni corrected).

Again, there was no obvious relationship between self-classified imagery group and scores for seen imagery. Many Projectors selected low scores for scenarios involving seeing or projecting images, while many individuals in the Inside and Off-screen groups reported seeing imagery often. Only the Verbalisers overwhelmingly endorsed the lowest imagery options.

#### Thought modality

We also asked participants to decide what the typical modality of their thoughts was. A narrow majority (50.9%) responded that they were mostly visual thinkers, while 27.8% endorsed the view that they thought with their inner voice but without pictures, 9.1% responded they think in words but without an inner voice, and 12.2% endorsed none of these options. The self-classified groups differed significantly in these responses (χ^2^=9.9, p=0.0194). Most individuals in the three imagery groups responded that they were visual thinkers, with the second largest proportion responding that they used their inner voice. Less than 14% in these three groups responded with either of the other options (Figure 5C). By contrast, most Verbalisers responded that they used their inner voice or thought in voiceless words, while 16.7% felt none of these options were applicable (Figure 5C, black bars). However, one Verbaliser (8.3%) considered themselves a visual thinker.

## Discussion

We sought to quantify how people experience their mental visual imagery. We found that commonly used vividness scores correlated widely with diverse measures of the strength of imagery and the propensity to visualise, as well as with measures of the richness and detail of automatically evoked pictorial representations. Importantly, we also quantified the experience of literally “seeing” imagery or projecting these experiences into the external world. Specifically, we asked to visualise an apple – either with eyes closed or with open eyes projected inside a frame on the screen. Our data suggest that people with vivid imagery are more likely to see or to project their imagery. Therefore, seeing, projecting, and vividness are *not* orthogonal dimensions. Crucially, seeing imagery is not only an extreme experience of hyperphantasic people. Rather, the clarity or intensity of seen imagery varied across the whole range. Many people with weak general vividness scores reported experiencing faint imagery that seemed to be projected externally, or that they could see faint images with their eyes shut.

Conversely, many individuals with high general vividness scores reported not seeing any image of an apple or only seeing a faint scheme of it. “Seeing” imagery therefore does not scale linearly with imagery vividness. Seeing imagery might trade off against self-reports of vividness due to the lack of an objective reference (Figure 1D,E). Someone who literally sees a faint image might rate this as not being very vivid. By contrast, someone who only reports the detail of their image within their metaphorical/internal mind’s eye might rate this as highly vivid.

### Vividness as measure of imagery

Nevertheless, vividness correlates strongly with many experiences. When asked about the appearance of an object that was given a high vividness score, people were able to report on its appearance (like whether a banana had black spots) and they volunteered extraneous visual details on free-text questions. People also overwhelmingly reported using imagery to compare the colour of unseen objects and showed a strong tendency to consider themselves visual thinkers. This appears to be independent of whether people experienced their mental images as an unseen internal representation, “seen” by the eyes, or even projected into the environment. By contrast, people with low vividness – especially those who identified as thinking about objects as verbal lists of facts – often found that these questions made no sense to them. Taken together, these findings suggest that vividness questionnaires can track the quality (whatever exactly that means) of mental pictorial representations. Vividness appears to be a robust feature of human cognition, even if it is hard to describe.

Critically, we found that the commonly used VVIQ-2 instrument, for quantifying vividness, may be redundant. A single 5-point cartoon version of a vividness questionnaire correlated more strongly with overall VVIQ-2 scores than most of the 32 items in that questionnaire correlated with one another. Both scores also revealed similar correlations with most other questionnaire items. For aphantasics this redundancy of individual questions in the VVIQ-2 can entrain a habit of responding with the minimum option – leading to inattention. Our attention checks suggested this could indeed have been the case: one attention check presented the same cartoon depictions of the apple as our imagery question, but instructed participants to respond with the maximum, the reddest apple. Three of the five individuals who failed this attention check had VVIQ-2 scores of 34 or less. The answers of these participants to other questions, especially the ones with free-text answers, did not suggest that they were generally disengaged from the task. But of course, we cannot know why any participants failed attention checks. Any participants who failed these checks were removed from analyses. Nevertheless, to reduce this potential problem – and to save time – we omitted the VVIQ-2 in our second survey. We generally replicated the relationships observed in the first survey with our simple cartoon question regarding the clarity of an imagined beach, although we note that the correlations in Experiment 2 using beach clarity as a proxy vividness were lower than those for the VVIQ-2 scores in Experiment 1. This could be because the demographics differed between the two experiments (see below).

While highly redundant, other evidence has suggested that if people are repeatedly asked to attempt to visualise, outcomes can be variable – with an imagined experience only conjured on a subset of trials [11,30]. So, while the repeated similar questions of the VVIQ-2 may be redundant to the point of disadvantage, they might also to some degree measure how reliably different people are able to conjure an imagined experience. However, the design of the VVIQ confounds trial-by-trial variability with different scenarios, which are presented in a fixed order. A direct test of fluctuations in imagery strength would be better addressed via a targeted investigation or instrument for this purpose. Moreover, it is also plausible that many people – especially those with intermediate levels of imagery – simply vary their responses from trial to trial at random to avoid responding with the same option every time. This variability would only constitute noise, reducing internal consistency.

Others researchers have described “objective” measures of imagery strength, such as perceptual priming of binocular rivalry [27,31–33] or pupil dilation [34]. These psychophysical and physiological measures correlate strongly with vividness ratings. However, vividness ratings are inherently subjective and therefore this is based on a circular logic: insofar as such measures replicate [35], these novel measures cannot be more objective than the self-reports used to identify them. What is more, they are contaminated by measurement noise and interindividual variability inherent in any experiment.

Therefore, the simplest way to study the subjective experience of mental imagery it to ask participants what they experience. We demonstrated that this can be achieved via a brief questionnaire, obviating the need for a protracted questionnaire. But of course, having only a single vividness question may not be optimal either. Future research on imagery could combine a small number of diverse questions, for example, our cartoon question, the question about imagining fiction, and a few selected VVIQ items. Ideally, items should also use inverted scales to minimise habitual responding.

Moreover, we certainly agree that research should relate results from subjective self-reports to perceptual or physiological experiments to investigate the neural basis of imagery. But such efforts can only seek to understand how mental imagery works, not replace self-reports.

### Self-classification into imagery types

In one question, we asked participants to imagine an apple with open eyes. In a follow-up question, we asked them to choose between four cartoons to indicate which had best illustrated what happened in their mind. One group, Verbalisers, chose an illustration of a thought bubble containing a list of facts. This is comparable to the lowest option in vividness questionnaires, such as our beach clarity question or the VVIQ. As such, it is unsurprising that Verbalisers consistently scored low on vividness and on measures of seeing imagery.

Interestingly, in Experiment 1 we also found that average vividness was somewhat reduced for Off-screeners compared to other imagery groups. Off-screeners chose an illustration with a thought bubble behind the head containing an apple. This could be considered as akin to “only knowing you are thinking of the object” rather than seeing it in the mind’s eye. This is the lowest vividness option in the VVIQ-2. However, we did not replicate this difference in Experiment 2 where Projectors, who chose the illustration of seeing an apple before them, scored lower on vividness than Off-screeners. Note that Experiment 2 is probably more representative of the general population because recruitment did not target communities interested in aphantasia or mental imagery research.

Numerically, Insiders, who identified with the illustration of holding a mental image inside the head, rated their imagery as more vivid than Projectors. In fact, Insiders were the only group that included participants with vividness scores greater than 145, which one might consider hyperphantasic. Taken together, this implies that Projectors are not characterised by more vivid imagery. It is possible that the act of projection weakens people’s impressions of imagery, as imagined images are either contrasted with actual visual inputs, or become intermixed with impressions of the external world. Imagined images that remain distinct from people’s impressions of the external world might be experienced as more vivid.

Curiously, all four groups included participants with very low vividness scores. A substantial number of Projectors had VVIQ-2 scores below 40 – some even scored the minimum of 32. This seems counterintuitive: many studies would categorise such individuals as aphantasic – the cut-off used to define aphantasia in many studies is considerably *higher* than 40 [1,27–30]. Why would anyone say that they have a mental image in front of them, but also respond like aphantasics in vividness questionnaires?

There could be several reasons. Of course, we cannot rule out that some people simply made a mistake in their self-classification. This could have been the case for two Projectors with minimal VVIQ-2, but it is unlikely to explain *all* the Projectors with low vividness scores. Importantly, the self-classification involved a single imagery scenario that was not designed to test participants’ *typical* experience of imagery. Therefore, some people may have experienced seeing an image of an apple on that particular trial but did not consistently form mental images across other questions.

Vividness questions could differ in terms of response criteria. The VVIQ-2 requires participants to choose between several descriptions of imagery vividness. The apple question provided several cartoon depictions, and asked participants to choose the one that most closely matched their experience of the mental image. Participants might find that “vague and dim” (option 2 in VVIQ) or a faint outline of an apple did not adequately reflect their experience. They might therefore choose the minimum, even though they mentally pictured an apple.

The VVIQ-2 also explicitly instructs participants to close their eyes while imagining scenarios [17]. By contrast, our self-classification scenario explicitly instructed participants to keep their eyes open. Intuitively, one might assume that visualisations are stronger with closed eyes, removing external stimulation. While the intensity of seeing the apple with eyes shut was strongly correlated with seeing it projected onto a screen, this does not rule out the possibility that these experiences dissociate for some people. Indeed, anecdotal feedback from two self-classified Projectors (one being an author) suggests that imagery for them is easier with open eyes.

Moreover, participants also vary in their other mental sensations. Anecdotally, several visual aphantasics have described to us that they imagine objects in tactile terms (see also [36]). Some might have felt the apple in front of the head best illustrated their experience: even though they did not experience the apple *visually*, they might nevertheless have conceived of it in space before them.

Interestingly, one recent study explored the perceived location of mental imagery [11]. It is unclear how these measures align with our self-classification question: Off-screeners might experience their images as having an undefinable location, but they could equally respond that they only “know” that they are thinking of an object (the lowest VVIQ option). Similarly, Projectors might have experienced an apple as being in front of them, but the cartoon could also be interpreted as merely “seeing” it with the eyes. Insiders, on the other hand, might also “see” the apple but locate it inside their head because they know it is not in the external world. Of note, we found in Experiment 1 that Insiders scored highest on vividness, and they were the only group that included people who are typically considered as hyperphantasic.

Unfortunately, the study quantifying the location of mental images did not include a traditional measure of the subjective vividness of imagery [11]. The authors argued that because imagery was unreliable across trials (experienced on 50% or less of all trials by 31% of people in their sample) and because it fluctuated in reported strength across trials within their study, this had rendered traditional vividness questionnaires [16,17] invalid. This, however, omits the possibility that traditional vividness scores might have inadvertently been measuring these qualities.

We note that the design of the four options (and the images used to represent them) was informed by our own experiences. Our author team includes individuals from each of the four groups. Moreover, these options reflect years of anecdotal discussions with people about how they do or do not experience mental imagery. Nevertheless, we cannot rule out that none of the four options were a good match for some people. People probably also vary in how they interpret these images. For example, a Projector might choose the Insider option because they feel that the apple inside the head better represents their experience of a mental image even if they “see” it before their eyes. Future research should seek to better define how individuals with imagery can be subdivided. This endeavour must involve detailed thematic analysis and other qualitative methods.

### Seeing imagery, projectors, and prophantasia

Our self-classification question probed whether people naturally see mental images in front of them without explicit instructions to project it. But many Insiders and Off-screeners also frequently experience imagery projected into the environment. Conversely, many Projectors scored low on our measures of projection. Therefore, while some people indeed report projecting images often, many people do not. They are *capable* of projecting mental images, but they do not always do it and they evidently do not do this automatically.

Previous research has coined the term “prophantasia” to describe the ability to project mental images externally [13,22]. This better captures the prevalence of this experience than merely asking how often people do this. It also aligns with anecdotal descriptions from people who told us that their mental images are “normally inside” but that they can also project them if they need or want to. It could be possible that people experience their internal imagery as strong and detailed, but their projections as faint or vague. Future research should test prophantasia by testing both the intensity/clarity of “seen” images and the sense of its location, that is, whether images are merely seen (but feel inside the head or in the eyes) versus appearing as located in the external environment.

The experience of seeing versus not-seeing mental imagery parallels reported distinctions in grapheme-colour synaesthesia [37]: projector synaesthetes report literally experiencing black letters as coloured. By contrast, associator synaesthetes only “know” that the letter has a colour without literally seeing that colour [38], and this distinction is correlated with brain structure [39]. We previously repurposed the same terminology to describe the distinction between people who see mental imagery and those who do not [12]. Intriguingly, synaesthetes with low or non-vivid imagery are more likely to be associators [40]. Future research should investigate these relationships in greater detail.

### Are people hallucinating?

For those of us who never see mental imagery, the whole concept of prophantasia is bewildering. Could it be true that a considerable proportion of people can wilfully conjure phantoms that they experience as being projected into their impressions of the external world? Do people really see pictures when they visualise something before their closed eyes? Questions like these are debated regularly when discussing mental imagery. Our surveys attempted to probe this specifically, by explicitly framing questions about whether people “literally see” images. Nevertheless, we cannot rule out that people took this less literally than we had intended. We believe several points speak against this possibility:

First, as discussed in the introduction, common descriptions in the imagery literature suggest that people do indeed have these experiences (Figure 1A-C), unfathomable as it might seem to some of us. Second, many visual experiences occur without physical stimulation, such as visual dreams, near-sleep (hypnagogic or hypnopompic) visual experiences, or the Tetris effect [24]. These experiences feel distinctly visual. Hallucinations are a well-known symptom of several pathologies. The neural mechanisms evidently exist to conjure phantom images that can appear atop an observer’s real visual input. These mechanisms could also operate in neurologically and psychiatrically healthy people. The difference between healthy prophantasia and pathological hallucinations could instead be a failure of reality monitoring [41–43].

Third, many of us who do not see mental images can experience other mental sensations, such as the experience of hearing an inner voice or having an imagined sensation of touch or smell. These experiences do seem comparable to how prophantasia apparently impinges into the visual sense. Thus, for some people the experience of seeing a mental visual image could be similar.

Research needs to devise better means of quantifying imagined experiences. Our scenarios asked participants which cartoons best matched their experiences of imagery. To calibrate these self-reports, one could also ask participants for a similar match for other internally generated visual experiences, like retinal afterimages, the Tetris effect [24], or the intensity of visual dreams. They could then use that as a reference against which to match their sensations of imagery.

### Vividness and aphantasia

Interestingly, our imagery vividness scores correlated with other internally generated visual experiences, like having visual dreams, near-sleep visions, or the Tetris effect. This extends previous reports that many aphantasics (as defined by vividness scores) report having visual dreams even though this is less common than for people in the normal vividness range [44]. This again suggests that whatever vividness scores measure, they tap into something consistent in human cognition, or at least in how they self-report a diverse range of internally-generated experiences.

It has been suggested that vividness measures introspective awareness of mental images [45]. Our results speak against that possibility. Individuals with low vividness scores – especially Verbalisers – generally endorsed views that their thoughts are encoded as semantic knowledge (possibly, but not necessarily, as mental vocalisations), rather than pictorially. These people also did not automatically evoke visual details of imagined objects. By contrast, most people who reported having visual imagery spontaneously had answers to these questions. It seems unlikely that these differences are simply because a subset of people is unaware of their mental imagery. The more parsimonious explanation is that they simply do not think in pictures at all. Consistent with this, aphantasics (defined as having minimum vividness scores) do not show any implicit priming effects, unlike hypophantasic individuals with low vividness scores [46].

Our data suggest that imagery vividness ratings are a robust measure, that accurately described people’s experiences of thinking. These data highlight important differences between groups of people with putatively similar imagery vividness scores. Any study of the cognitive or neural mechanisms of imagery that uses lax criteria, for instance to separate people with no imagery from people who can sometimes visualise, will likely conflate people with distinct imagery profiles and capacities. We therefore agree with others [46–48] that we need a stricter definition of aphantasia that excludes anyone who experiences mental images, however faint.

## Data availability

All materials, data, and analysis code are publicly available at https://osf.io/eu294

## Supplementary Results

### Lack of correlations with vividness

Vividness scores (VVIQ-2) in Experiment 1 did not predict how often participants felt depictions in movies or television affect the mental images created by reading fiction (ρ=0.3, p=0.8493). Beyond imagery, vividness also did not correlate with whether dreams feel real (ρ=0.2, p=1), and participants’ tendency to mind-wander (ρ=0.1, p=1) or be absentminded (ρ=0.1, p=1). While it was moderately correlated with participants’ tendency to daydream (ρ=0.39, p<0.026), it did not predict whether they consider their daydreams distracting (ρ=0.23, p=1).

### Daydreaming habits

One could assume that people with vivid imagery – but especially if projected externally into the environment – might be more absentminded, distracted, and have greater tendency to mind-wander. But as mentioned, the relationships between these variables and vividness scores were at best weak. However, our data revealed relationships between participants’ daydreaming habits.

Absentmindedness, mind-wandering, and daydreaming were all correlated (minimum: ρ=0.24, p=0.024). Individuals who daydreamed often also typically experienced their daydreams visually (ρ=0.67, p<0.0001). In turn, having visual daydreams correlated strongly with experiencing daydreams projected into the environment (ρ=0.72, p<0.0001).

### Qualitative analysis of free-text answers

Many individuals provided elaborate descriptions of the appearance of the elephant beyond the basic description of its pink colour. They often described the eyes and ears or the texture of the elephant’s skin, its size and pose, and sometimes even details of the surrounding scene. In one description, the elephant wore a harness. Similarly, for the triangle they were more likely to describe the colour, material, how the shape was drawn, extraneous details not implied by the question, or the surrounding space. Several people described the musical instrument.

However, low-vividness individuals (VVIQ-2≤40) mostly restricted themselves to descriptions that did not go beyond restating the instruction. For the elephant question, example responses were “pink and big,” “The elephant is pink,” and “A standard elephant which is otherwise pink in colour”, or generic descriptions of elephants that did not imply any specific image. One individual with VVIQ-2 of 35 stated “my brain goes: pink elephant, pink elephant, pink elephant…” Similarly, their responses to the triangle question were often generic descriptions of what a triangle is, such as “3 edges and 3 vertices,” “3 straight sides, 3 angles,” and “A shape with 3 sides.” However, consistent with the lower correlations between the ratings of the triangle and vividness, this pattern was less clear. One individual with VVIQ-2 of 113 simply responded with “a^2^+b^2^=c^2^”. Two individuals with VVIQ-2 of 105 and 158, respectively, did not enter any description of the triangle.

Interestingly, for many participants the triangle was either equilateral or an isosceles triangle. Across the whole sample, 32.6% of responses contained the words “equilateral”, “equal”, or “same”. These proportions were similar for low vividness (VVIQ-2≤40) participants compared to the others (30% vs 33.3%, respectively).

### Triangle question in Experiment 2

We instructed participants to think of a triangle and then asked them to report its colour. This divided the sample roughly in three equal sets: 39.6% responded that they knew the colour, 30.9% responded that they did not, and 29.6% responded that this question made no sense. These groups did not differ in vividness scores (χ^2^=0.21, p=0.9009).

### Familiarity with aphantasia

We quantified how familiar participants were with aphantasia. We asked them to choose between three options describing what an aphantasic person experiences when they think of an object. Most participants responded they did not know or had never heard of aphantasia (48.3%), 36.5% responded that aphantasics think in terms of facts but have no mental images, while 15.2% responded that aphantasia is the absence of “literally seeing” an object.

---

**Figure S1.**
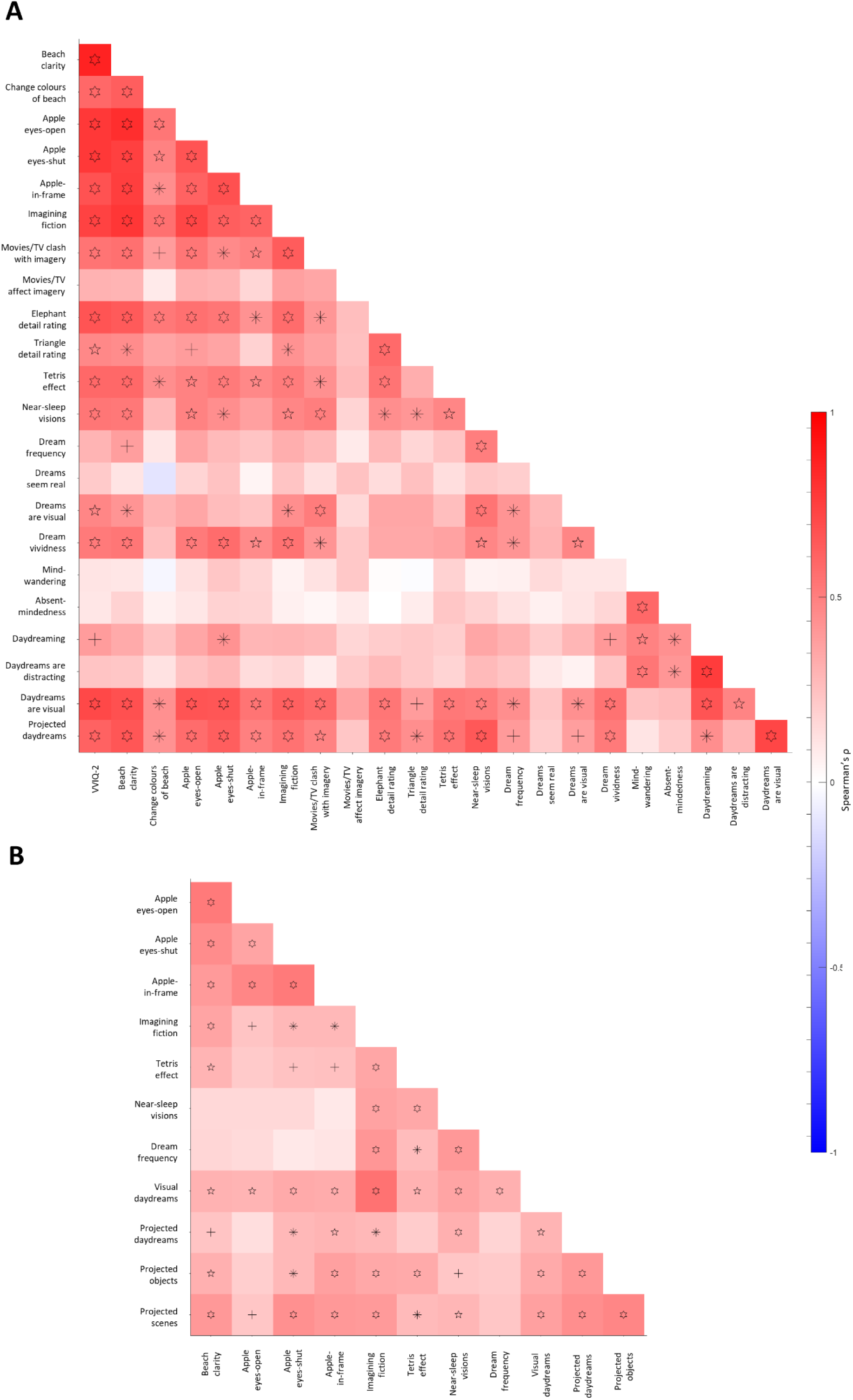
Correlation matrix (Spearman’s ρ) between survey items with Likert-scale responses in Experiment 1 (**A**) and Experiment 2 (**B**). Symbols denote statistical significance after Bonferroni correction. Plus sign: p<0.05. Asterisk: p<0.01. Pentagram: p<0.001. Hexagram: p<0.0001.

**Figure S2.**
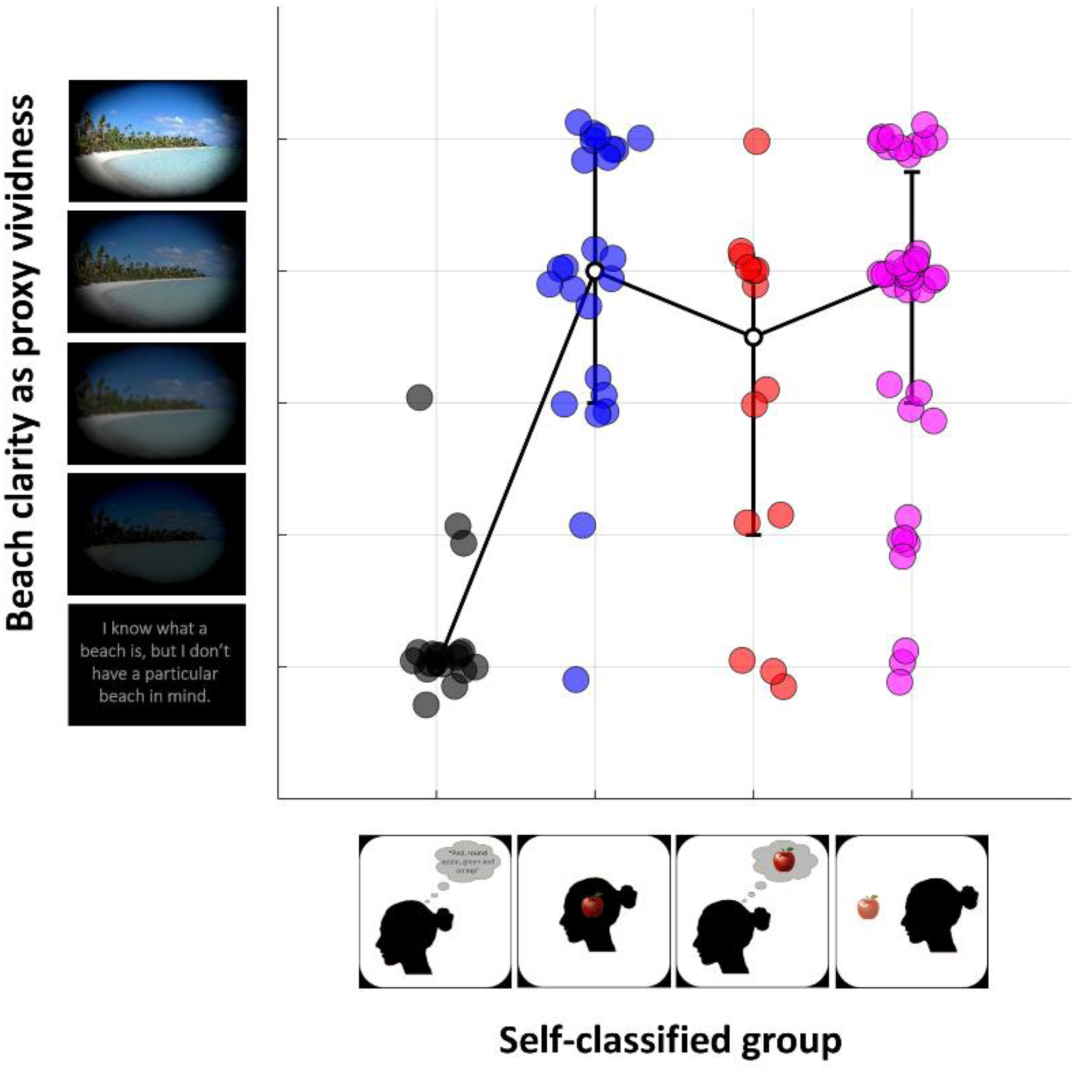
Beach clarity scores for each group from self-classification in Experiment 1. Each dot is one participant. Some Gaussian jitter was added to dot locations for visualisation. The solid black lines connect the medians (error bars denote the interquartile range). Black: Verbalisers. Blue: Insiders. Red: Off-screeners. Purple: Projectors. Beach clarity scores differed significantly between groups (Kruskal-Wallis test, χ^2^=38.4, p<10^−7^, uncorrected), with Verbalisers scoring significantly lower than the imagery groups (Mann-Whitney U-test, all p<0.001). Off-screeners scored lower than Insiders (p=0.0311) but were not significantly from Projectors (p=0.0897). Insiders and Projectors were not significantly different (p=0.3844).

**Figure S3.**
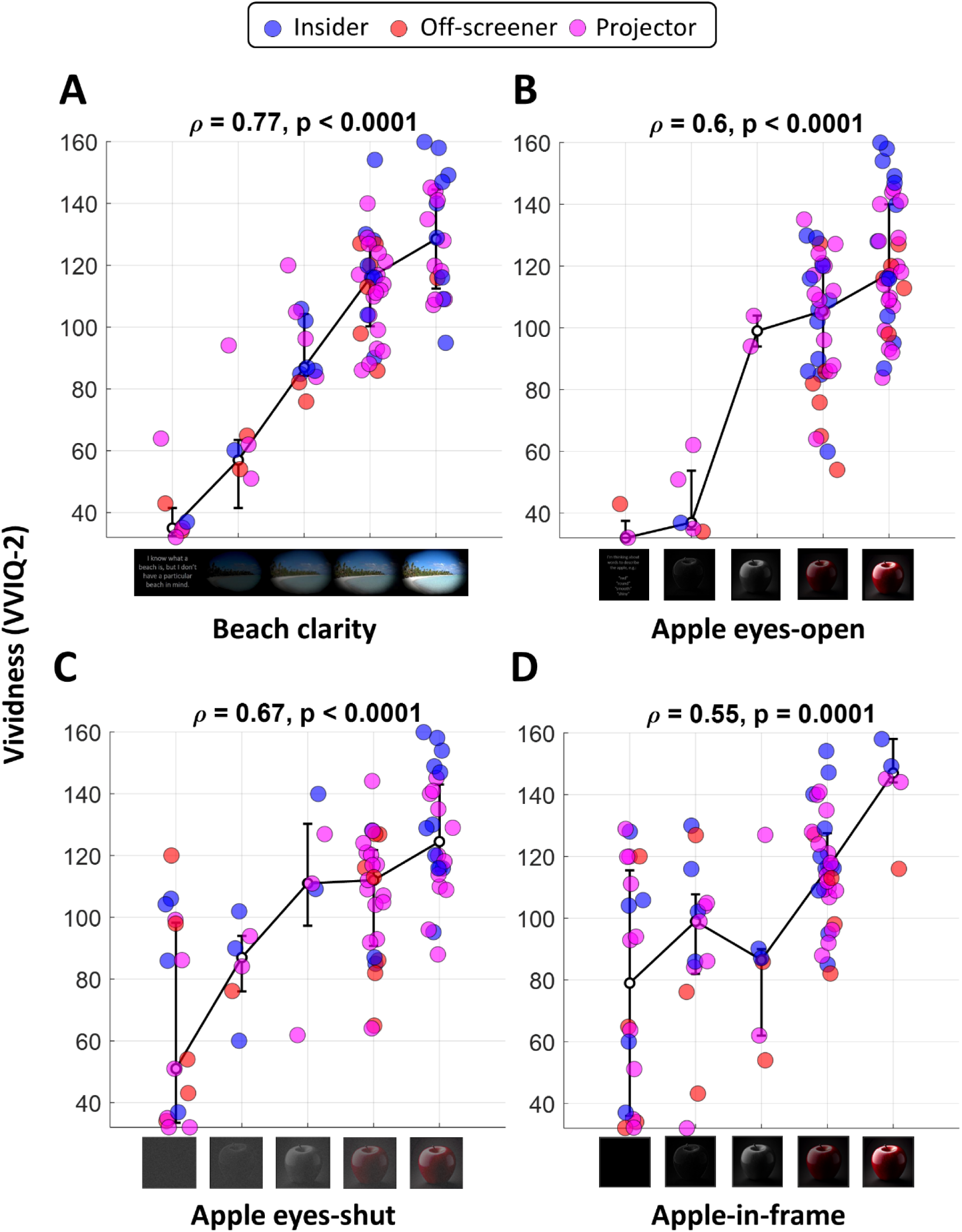
Vividness (VVIQ-2) scores from Experiment 1, excluding the Verbaliser group. Data are plotted against the Likert score on the beach clarity (**A**), apple eyes-open (**B**), apple eyes-shut (**C**), apple-in-frame (**D**) scenarios. Each dot is one participant. Some Gaussian jitter was added to dot locations for visualisation. The solid black lines connect the medians (error bars denote the interquartile range). Blue: Insiders. Red: Off-screeners. Purple: Projectors. Statistics at the top of each panel show the Spearman correlation (Bonferroni corrected).

